# Distinct Metabolic Signatures Linked to High-Resolution Computed Tomography Radiographic Phenotypes in Stable and Progressive Fibrotic Lung Disease

**DOI:** 10.1101/2025.10.30.685696

**Authors:** Girish B. Nair, Faizan Faizee, Zachary Smith, Sayf Al-Katib, Nadia Ashrafi, Ali Yilmaz, Romana Ashrafi Mimi, Vilija Lomeikaite, Juozas Gordevičius, Edward Castillo, Stewart F. Graham

## Abstract

**Rationale and Objectives:** This study aimed to identify distinct metabolic signatures associated with disease progression by integrating high-resolution computed tomography (HRCT) visual scoring with comprehensive metabolomic profiling.

**Materials and Methods:** This single-center, cross-sectional study enrolled 60 idiopathic pulmonary fibrosis/interstitial lung disease (IPF/ILD) patients from December 2021 to October 2022. Participants underwent standardized pulmonary function testing, HRCT imaging, and peripheral blood collection for metabolomic analysis using one-dimensional hydrogen nuclear magnetic resonance (^1^H NMR) spectroscopy and ultra-high-performance liquid chromatography coupled to tandem mass spectrometry (UPLC-MS/MS). A comprehensive HRCT scoring system quantified total HRCT composite scores. Linear regression analysis integrated radiographic scores with metabolomic profiles, adjusted for multiple covariates

**Results:** Stable IPF/ILD exhibited moderate negative correlations between the six most significant metabolites and HRCT scores (r = −0.27 to −0.51), along with a high abundance of specific phospholipids (triacylglycerol, monoacylglycerol, phosphatidylglycerol, phosphatidylethanolamine, diacylglycerol), sphingomyelin, ceramide, and acylcarnitine. In contrast, progressive disease showed weak positive correlations between the six most significant metabolites and HRCT scores (r = 0.19–0.26), and moderate negative correlation between specific triacylglycerol species and HRCT scores (r = −0.37-0.4). Furthermore, metabolomic analysis in individuals with progressive disease revealed both high and low abundances of specific phospholipid species (including high and low triacylglycerol species, as well as low levels of phosphatidylglycerol, phosphatidylethanolamine, phosphatidylcholine, phosphatidylserine, and phosphatidylinositol), along with high levels of certain sphingomyelin, ceramide, taurine, and purine bases, and low levels of xanthine and lactic acid observed.

**Conclusion:** Integration of systematic HRCT visual scoring with metabolomic profiling successfully differentiated stable from progressive IPF/ILD through distinct molecular-radiographic signatures.

**Funding:** Supporting Effective Educator Development (SEED) grant

## Introduction

Fibrotic interstitial lung disease (F-ILD) encompasses a heterogeneous group of progressive respiratory disorders characterized by pathological extracellular matrix deposition and disruption of normal lung parenchymal architecture. While idiopathic pulmonary fibrosis (IPF) represents the archetypal and most severe phenotype among F-ILDs, the distinction between stable and progressive disease trajectories has emerged as a critical prognostic determinant. Progressive FILD is distinguished by recurrent acute exacerbations, markedly reduced survival, and substantially increased healthcare utilization and economic burden.^1^

High-resolution computed tomography (HRCT) plays a crucial role in diagnosis, with specific attention to the presence of usual interstitial pneumonia (UIP) patterns.^2^ This imaging modality enables detailed characterization of pulmonary parenchymal architecture and associated pathological features. HRCT enables detailed evaluation of pulmonary fibrosis through both qualitative and quantitative methodologies. Conventional visual scoring, based on radiologist interpretation, and automated quantitative assessment are widely employed to assess the extent of fibrosis. The prognostic utility of both scoring systems is well-established, with evidence demonstrating their value as independent predictors of disease outcomes^3–9^. Furthermore, metabolomics, the comprehensive study of samples (metabolites) within biological systems, has identified distinct metabolic signatures associated with interstitial lung disease progression. These findings suggest that metabolomic profiling could complement imaging-based metrics to enhance prognostication and refine disease stratification.

In this cross-sectional study, we investigated the correlation between plasma metabolomic profiles and HRCT visual scoring patterns in both stable and progressive F-ILD/IPF cohorts. We hypothesized that specific metabolic signatures would demonstrate significant associations with the extent of radiographic abnormalities, potentially offering novel insights into disease mechanisms and progression. Through multivariate linear regression analysis, we examined the relationship between key metabolites and composite HRCT visual scores, while adjusting for relevant clinical and demographic variables. This investigation represents a crucial step toward integrating radiographic findings with metabolic pathways for enhanced disease characterization/classification and disease progression in fibrotic lung disease.

## 2.0 Methods and Materials

### 2.1 Study Design and Patient Population

This single-center, cross-sectional study was conducted from December 2021 to October 2022 following approval from the Corewell Health Institutional Review Board (IRB 2021-327) and supported by a Supporting Effective Educator Development (SEED) grant. Of the 196 screened patients with confirmed diagnoses of ILD/IPF, 60 participants met the predetermined inclusion criteria (Supplemental Figure 1). All participants underwent standardized pulmonary function testing, a high-resolution CT scan without IV contrast (HRCT), and peripheral blood sample collection for metabolomic profiling within three months of enrollment. Participant stratification into progressive or stable ILD/IPF cohorts was based on retrospective assessment of clinical trajectory over the 12 months preceding enrollment, adhering to the 2018 IPF and Progressive Pulmonary Fibrosis (PPF) guidelines^10^. Disease progression was defined by the presence of at least two of the following criteria within the preceding 12 months: (1) absolute decline in forced vital capacity >10% or diffusion capacity >15%, (2) radiographic progression involving >10% of lung parenchyma, or (3) documented clinical deterioration of respiratory symptoms. Comprehensive baseline demographic and clinical characteristics were systematically recorded, including age, race, gender, BMI, tobacco use disorder, family history of lung cancer, relevant medical comorbidities, pulmonary function test (PFT), and current antifibrotic or immunosuppressive therapeutic regimens (Supplemental Table 1 and 1B).

### 2.2 Proton Nuclear Magnetic Resonance Spectroscopy (^1^H NMR) Sample Analysis and Data Acquisition

Blood samples were collected after an overnight fast (minimum of 8 hours) to minimize the influence of dietary factors on lipid profiles. Collection was performed using standard venipuncture techniques with serum separator tubes. Serum was separated by centrifugation at 1,200 *g* at 4°C for 10 minutes. Serum was aliquoted into cryovials and stored at −80°C until analysis. All samples were processed within 2 hours of collection to maintain metabolite stability.

Serum samples (300 µl) were filtered through pre-rinsed (washed with water x7 at 4°C and centrifuged at 12,000 *g* for xxx mins) 3.5 KDa filters via centrifugation (13,000 *g*, 4°C, 30 min) to remove proteins and macromolecules potentially interfering with the spectral baseline. The filtrate (228 µl) was combined with D_2_O (28 µl) and DSS (24 µl of 11.77 mM sodium 2,2-dimethyl-2-2silapentane-5-sulfonate in 50-mmol NaH_2_PO_4_ buffer). Samples were analyzed at 310 K using a Bruker Ascend III HD 600.13 MHz spectrometer with a TCI probe. 1D ^1^H-NMR spectra were acquired using noesygppr1d pulse sequence (256 FIDs, 64 K complex points, 20 ppm spectral window, 5-second relaxation delay). Metabolite identification and quantification were performed using Bayesil^11^.

### 2.3 Liquid Chromatography-Mass Spectrometry-based Sample Analysis and Data Acquisition

For liquid chromatography and mass spectrometry (LC-MS) analysis, high-purity LC-MS grade solvents (water, acetonitrile, methanol, isopropyl alcohol, formic acid) were obtained from Fisher Scientific (Hanover Park, IL, USA), and ethanol, pyridine, and phenylisothiocyanate were from purchased from Sigma Aldrich (St. Louis, MO, USA).10 µL of the serum aliquot was prepared as per the manufacturer’s instructions (biocrates Life Sciences, AG, Innsbruck, Austria). Briefly, sample preparation included nitrogen drying, phenylisothiocyanate (PITC) derivatization, and extraction in 5 mM ammonium acetate in methanol. Extracts were analyzed using a Waters I-class Ultra Performance Liquid Chromatography (UPLC) unit coupled with a Sciex 7500 mass spectrometer (AB Sciex LLC, Framingham, MA, USA). The Q500 XL kit provided direct flow injection analysis (FIA) and FIA in extended load (FIA XL) for lipid analysis. The Q500 XL kit included three quality control (QC) levels: (a) QC1 (Low): Low concentration standards for each metabolite class; (b) QC2 (Mid): Medium concentration standards used for result normalization; and (c) QC3 (High): High concentration standards for linearity assessment. QC2 was used for result normalization. QC samples were analyzed at the beginning, middle, and end of each analytical batch to monitor instrument performance and ensure data quality. MetaboINDICATOR™, integrated into the WebIDQ™ software from biocrates, to calculate concentration, metabolite sums, and ratios for enhanced analysis of metabolomic and lipidomic datasets. We utilized the results from these metabolite indicator ratio calculations.

### 2.4 HRCT Scoring System Development and Implementation

All HRCT examinations were independently evaluated by the thoracic radiologist (S.A.) with over 10 years of experience in interstitial lung disease imaging, who was blinded to clinical data. The scoring system evaluated four cardinal radiographic features: ground-glass opacity (GGO), reticulation, traction bronchiectasis, and honeycombing. Each lung was systematically divided into three zones, yielding six total zones (R1-R3 and L1-L3): (1) Upper zone (R1/L1): apex to carina, (2) Middle zone (R2/L2): carina to inferior pulmonary vein, and (3) Lower zone (R3/L3): inferior pulmonary vein to base. Within each zone, features were scored on a semi-quantitative scale: (0) Absent, (1) Minimal (<5% involvement), (3) Mild (5-25%), (4) Moderate (26-50%), (5) Severe (51-75%), and (6) Very severe (>75%). This systematic approach generated multiple analytical parameters: (1) Individual feature scores, (2) Zonal scores, (3) Cumulative feature scores across zones, (4) Composite zonal scores, and (5) Composite HRCT scores.

### 2.5 Statistical Analysis

Statistical analyses were performed using R (v3.50.3) to compare demographic characteristics, clinical parameters, HRCT scores, and metabolite concentrations between different groups. Differential metabolite analysis was conducted using linear regression models. The analysis examined correlations between metabolite levels and HRCT composite scores within each disease group separately, adjusting for relevant covariates. Descriptive statistics, including medians and interquartile ranges (IQR), were calculated for each radiological feature. Between-group comparisons were performed using the Wilcoxon rank-sum. Parameters with significant differences are annotated in the figure, and detailed results are summarized in Supplemental Tables 2 and 3. Differential metabolite analysis was conducted using linear regression models implemented through the R package limma (version 3.50.3). The analysis examined correlations between metabolite levels and HRCT composite scores within each disease group separately, adjusting for relevant covariates. Metabolite concentrations were log2-transformed and normalized using quantile normalization to ensure distributional assumptions were met. The linear regression model was adjusted for several covariates, including HRCT visual scores (composite scores), age, gender, tobacco pack years, BMI, and use of antifibrotic and immunosuppressive therapies. Models were fitted using robust regression with empirical Bayes moderation through the lmFit function (maximum iterations: 100). Empirical Bayes statistics were computed using the eBayes function with robust variance estimation. Multiple testing correction was performed using the Benjamini-Hochberg false discovery rate (FDR) method.

## 3.0 Results

### 3.1 Demographic and Clinical Characteristics

The median age of patients in the Progressive IPF/ILD disease group (PD) was 75 years, compared to 74 years in the Stable IPF/ILD disease group (SD). Most participants in both groups were self-reported as White (90.9% in PD vs. 92.6% in SD), with a smaller proportion identifying as Black (9.1% in PD vs. 7.4% in SD) (Supplemental table 1).

Males comprised 54.5% of the PD group compared to 29.6% of the SD group. The prevalence of chronic obstructive pulmonary disease (COPD) was comparable between the groups (15.2% in PD vs. 18.5% in SD). Coronary artery disease (CAD) was more common in the SD group (44.4%) compared to the PD group (30.3%). Similarly, type 2 diabetes mellitus (T2DM) was slightly more prevalent in the SD group (25.9%) compared to the PD group (21.2%). Antifibrotic therapy was more frequently used in the PD group (63.6%) compared to the SD group (44.4%). Immunosuppressive therapy usage was almost similar between the groups (27.3% in PD vs. 29.6% in SD). (Supplemental table 1). Further, pulmonary function test in progressive disease exhibited lower mean FVC and DLCO values compared to stable disease (Supplemental Table 1B).

### 3.2 Comparison of HRCT Composite Scores Between Progressive and Stable Disease Groups

The median and interquartile ranges (IQRs) for each radiological feature, stratified by disease state, are presented in Figure 1. Reticulation scores were significantly higher in the PD cohort compared to the SD cohort (median [IQR]: 18.0 [16.0–20.0] vs. 14.0 [10.5–17.5], p = 0.0008). Similarly, Traction Bronchiectasis scores were elevated in the Progressive group (14.0 [12.0–17.0] vs. 12.0 [7.0–13.5], p = 0.0129). Composite HRCT scores also demonstrated a significant difference, with higher scores observed in the PD cohort (57.0 [44.0–60.0] vs. 46.0 [26.0–53.0], p = 0.0112). No significant differences were observed for GGO or honeycombing scores; however, an upward trend of honeycombing scores was seen in the PD. These findings highlight the distinct radiological features associated with disease progression in IPF/ILD cohorts. Boxplots (Figure 1) illustrate the distribution of radiological features across the two groups, highlighting the significant differences in Reticulation, Traction Bronchiectasis, and Composite Scores.

**Figure 1.**
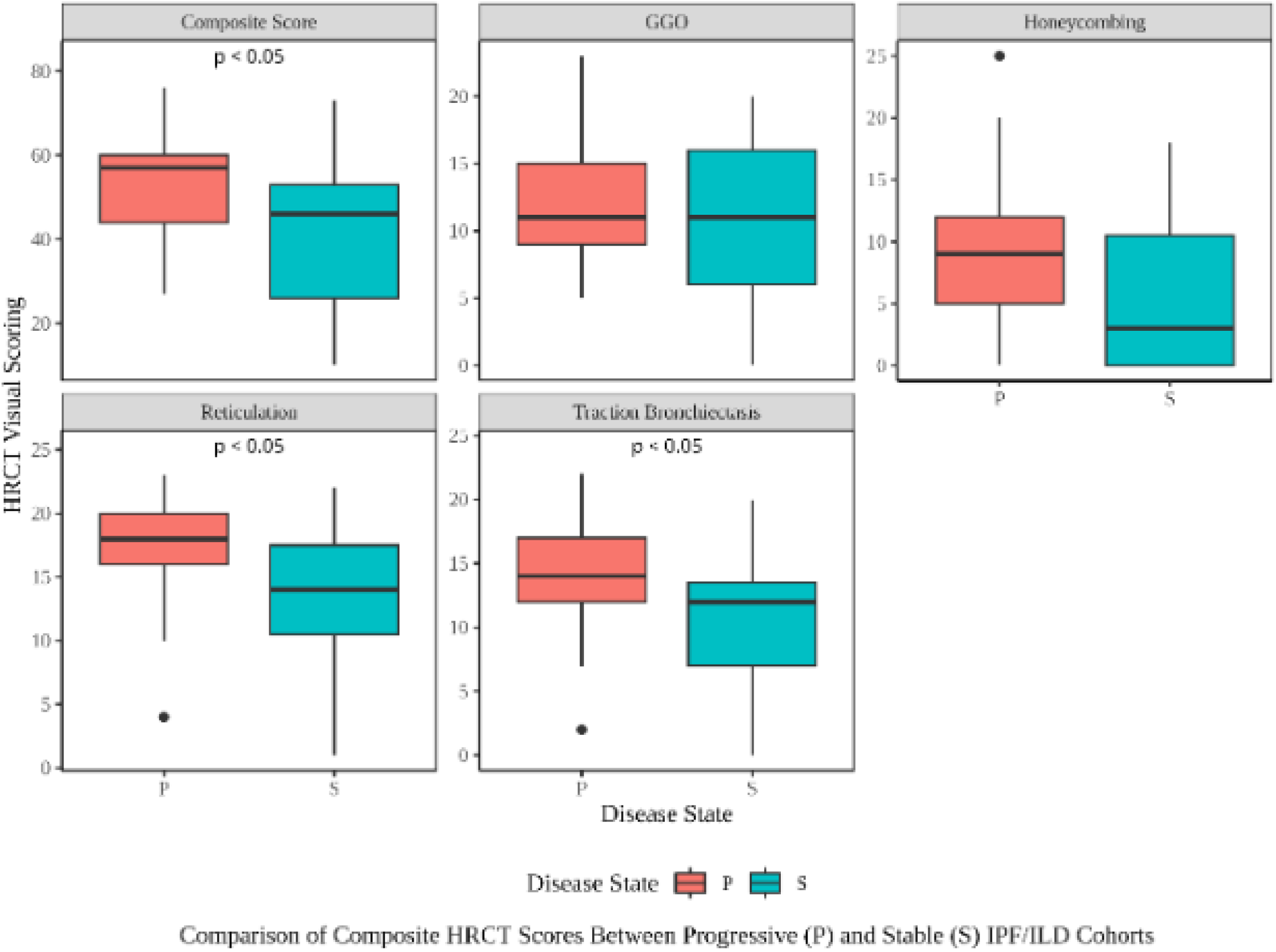
Comparison of HRCT Radiographic Features Between Progressive and Stable Disease Groups. Box plots comparing high-resolution computed tomography (HRCT) visual scoring features between progressive disease groups. Each box represents the interquartile range (IQR) with the horizontal line indicating the median value. Whiskers extend to the most extreme data points within 1.5 times the IQR, and individual points beyond the whiskers represent statistical outliers. Significant differences between P and S groups (p < 0.05) are indicated. Ground-glass opacity (GGO), reticulation, traction bronchiectasis, honeycombing, and composite scores were assessed. Reticulation, traction bronchiectasis, and composite scores demonstrated statistically significant differences between the two groups, with higher scores observed in the progressive disease group.

### 3.3 Metabolomic Correlations with Radiographic Feature in Stable and Progressive IPF/ILD

Linear regression analysis with multiple covariates, including the composite HRCT scoring of four radiographic patterns in stable and progressive disease cohorts, unveiled unique metabolic signatures (Figure 2 and Supplemental Tables 2A, 2A-1, 2A-2, 3A, 3A-1, 3A-2).

**Figure 2:**
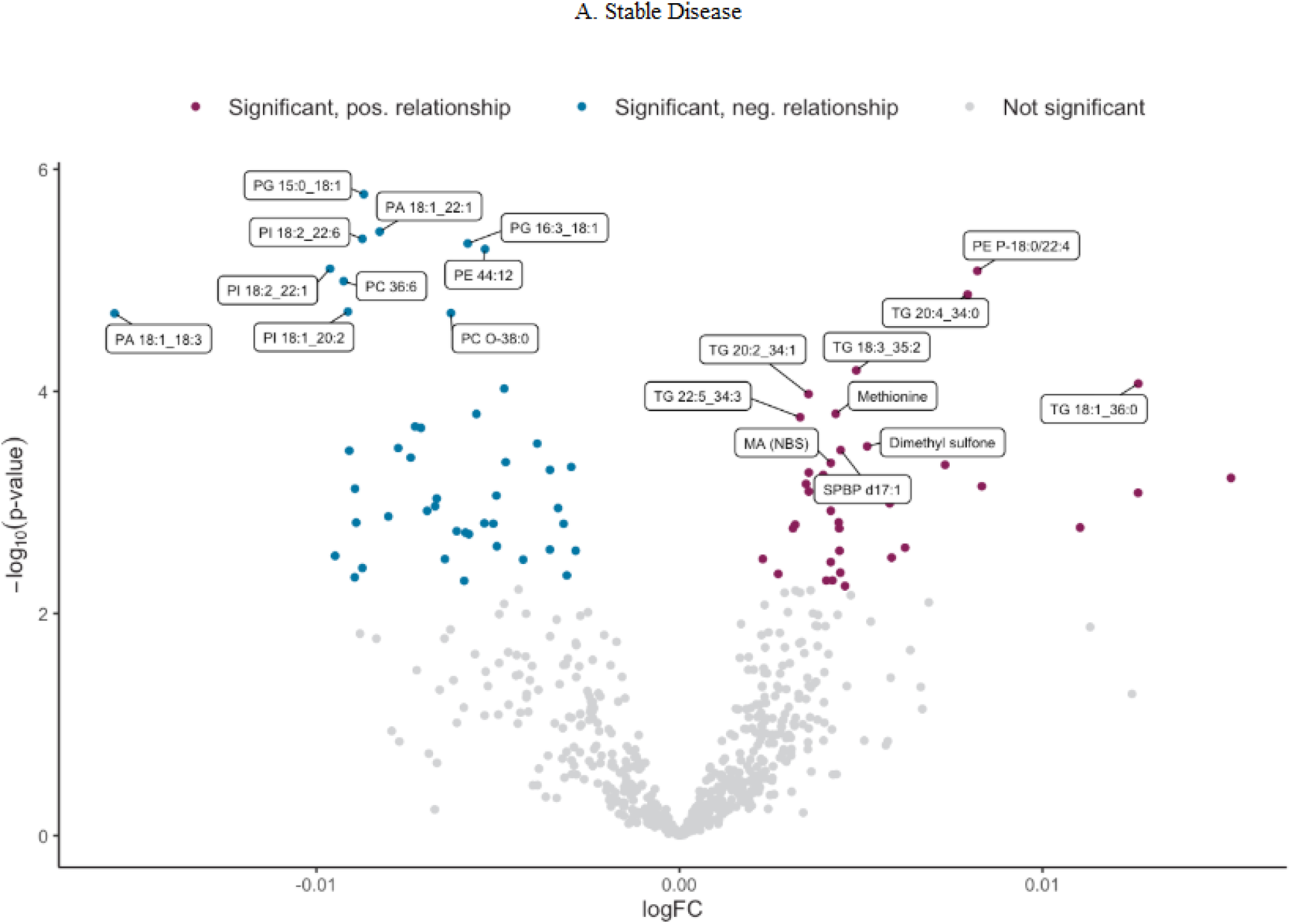
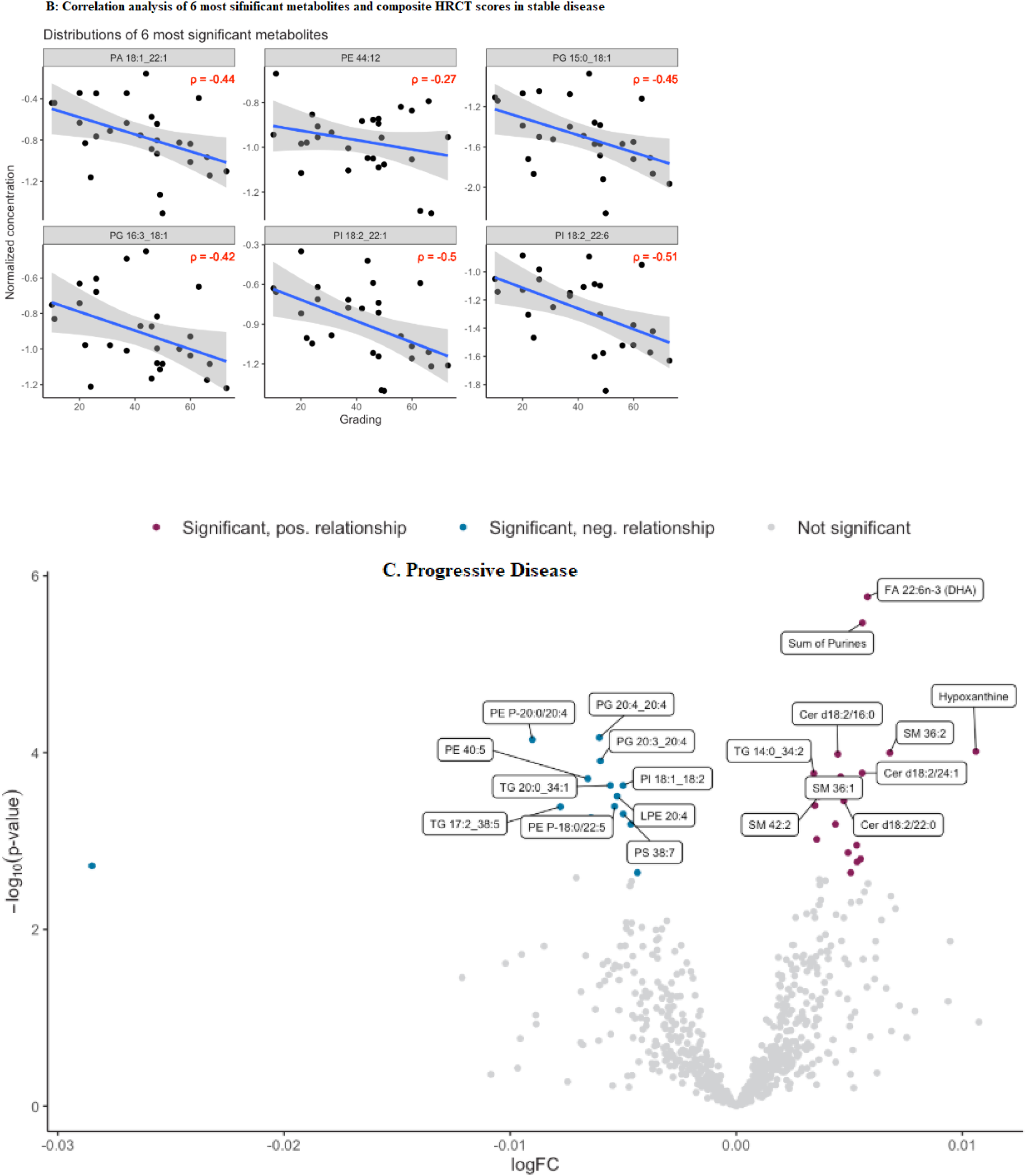
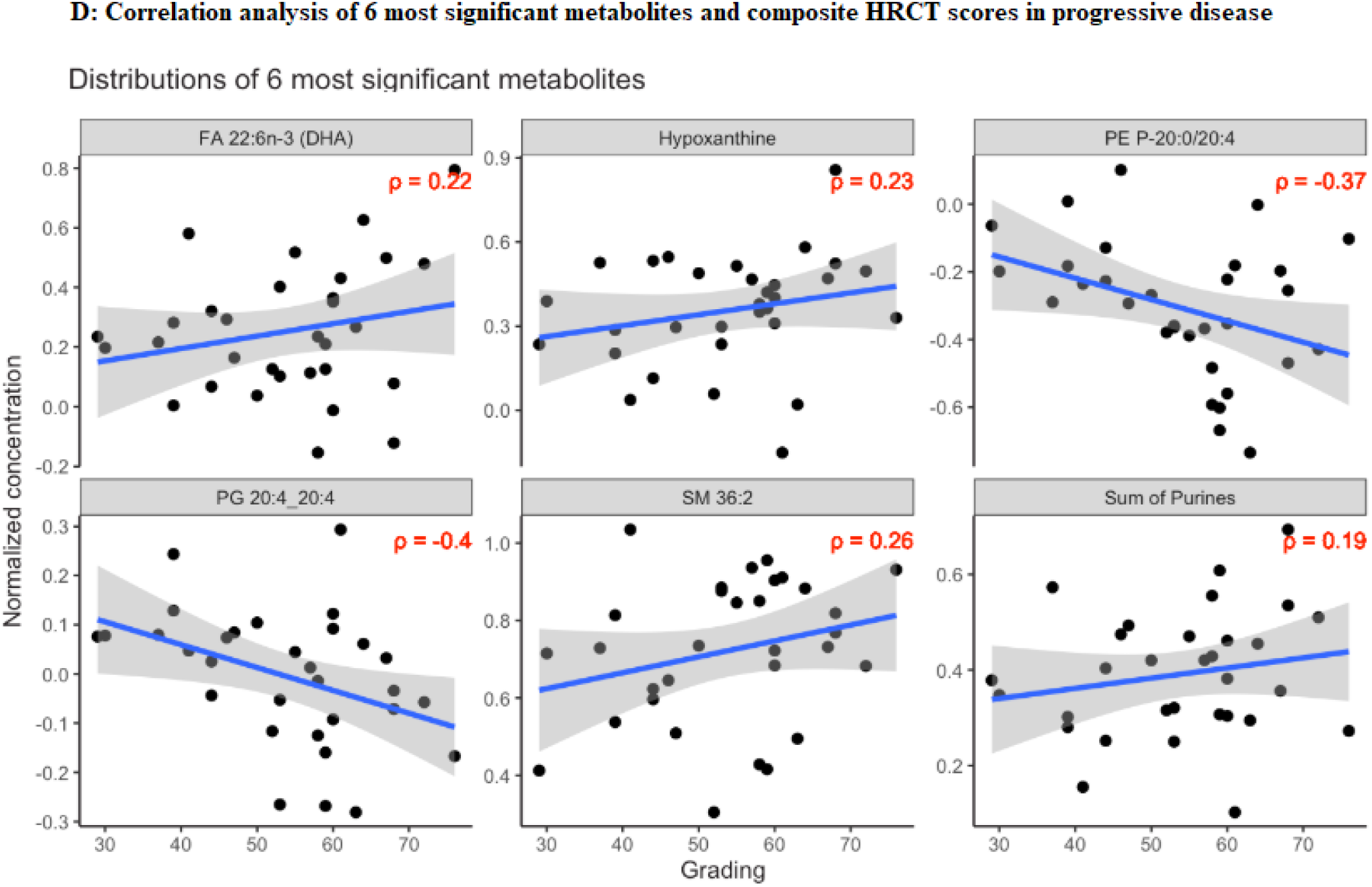
Figure 2A: Volcano plot of metabolite associations with stable disease. Scatter plot displays metabolites with log fold change (logFC) on the x-axis and –log (p-value) on the y-axis. Metabolites with significant positive relationships (burgundy) and significant negative relationships (teal) are highlighted, with selected metabolites labeled. Non-significant metabolites are shown in grey. **Figure 2C.** Volcano plot of metabolite associations with progressive disease. Scatter plot displays metabolites with log fold change (logFC) on the x-axis and −log(p-value) on the y-axis. Metabolites with significant positive relationships (burgundy) and significant negative relationships (teal) are highlighted, with selected metabolites labeled. Non-significant metabolites are shown in grey. **Figure 2B:** Correlation analysis of the six most significant metabolites with composite HRCT scores in stable disease. Each panel shows the normalized concentration of a metabolite (y-axis) and HRCT grading (x-axis). Pearson correlation coefficients (p) are indicated. **Figure 2D:** Correlation analysis of the six most significant metabolites with composite HRCT scores in progressive disease. Each panel shows the normalized concentration of a metabolite (y-axis) versus HRCT grading (x-axis). Pearson correlation coefficients (p) are indicated.

Linear regression analysis with HRCT composite scores in stable IPF/ILD participants revealed significant associations with several metabolites (FDR q-value < 0.05). High-abundant metabolites included several triacylglycerols, diacylglycerol species, monoacylglycerol, phosphatidylethanolamines, phosphatidylinositols, phosphatidylglycerols, sphingomyelin and ceramides, acylcarnitine, and betaine (FDR q <0.05; Supplemental Table 3A-1).

In contrast, stable ILD/ILD individuals revealed a significant decrease in the abundance of triacylglycerols, phosphatidylglycerols, phosphatidylethanolamines, phosphatidic acids, lactic acid (Lac), cholesterol ester, and indicators of beta oxidation metabolism (FDR q <0.05; Supplemental Table 3A-1).

Further, linear regression analysis with HRCT composite scores in progressive IPF/ILD individuals found several differentially significant metabolites (FDR q-value < 0.05). High abundant metabolites included triacylglycerols, a phosphatidylglycerol (PG 16:0_20:3), sphingomyelin and ceramides, hypoxanthine, a fatty acid (FA 22:6n:3 DHA), a cholesterol ester (CE 20.3), taurine, aspartic acid and the sum of purine metabolism (Supplemental Table 2A-1).

Conversely, low abundance metabolites included several triacylglycerols, phosphatidylglycerols, phosphatidylethanolamines, a phosphatidylcholine (LPC 20:4), a phosphatidylserine (PS 38:7), a phosphatidylinositol (PI 18:1_18:2), a bile acid (TMCA), and an indicator of xanthine synthesis metabolism (Supplemental Table 3A-2).

Finally, we performed a Pearson correlation analysis between the top six significantly abundant metabolites and total HRCT scores between progressive and stable ILD/IPF individuals. In stable IPF/ILD individuals, we found moderate negative correlation between significant low abundance metabolites and increasing total HRCT scores: PA 18:1_22:1 (r = −0.4), PE 44:12 (r = −0.27), PG 15:0_18:1 (r = −0.45), PG 16:3_18:1 (r = −0.42), PI 18:2_22:0 (r = −0.5), and PI 18:2_22:6 (r = −0.51) (Figure 2).

In progressive IPF/ILD individuals, we found a weak positive correlation between significant high abundant metabolites and total HRCT scores: FA 22:6n3 DHA (r = 0.22), hypoxanthine (r = 0.23), SM 36:2 (r = 0.26), sum of purine metabolism indicator (0.19). Additionally, a moderate negative correlation was observed between the significantly low-abundant metabolites and total HRCT scores: PE. P 20:0_20:4 (r = −.37) and PG 20:4_20:4 (r = −0.4).

## 4.0 Discussion

Our cross-sectional analysis revealed distinct metabolic reprogramming patterns that differentiate stable from progressive fibrotic lung disease through integrated HRCT and targeted metabolomics.

The most clinically significant finding demonstrates meaningful correlations between HRCT composite scores and specific metabolite classes in patients with IPF and other interstitial lung diseases. In progressive disease, we identified weak but statistically significant positive correlations between elevated metabolites and HRCT scores, including fatty acids (r = 0.22), hypoxanthine (r = 0.23), sphingomyelins (r = 0.26), and purine metabolism indicators (r = 0.19). Moreover, moderate negative correlations emerged between decreased phospholipid species and total HRCT scores across both disease states, with correlation coefficients ranging from −0.27 to −0.51. These correlations remained statistically robust after false discovery rate correction and adjusting for several covariates, supporting biologically relevant relationships between systemic metabolic alterations and the extent of structural lung injury quantified by HRCT. The significantly higher composite HRCT scores observed in progressive disease (median 57.0 vs. 46.0, p <0.05) were primarily driven by increased reticulation (18.0 vs. 14.0, p <0.05) and traction bronchiectasis scores (14.0 vs. 12.0, p <0.05). These radiographic differences correspond with distinct metabolic reprogramming patterns, suggesting that the structural features quantified on HRCT reflect underlying metabolic dysfunction that may precede or parallel the morphologic changes visible on imaging.

Both disease groups exhibited alterations across multiple lipid metabolite classes, though with distinct patterns. In stable IPF/ILD, lipid metabolism demonstrates a complex and dynamic profile characterized by extensive TG changes, with several TGs showing increased abundance alongside selective decreases in other TG species, phosphatidylglycerol, and phosphatidic acids. This complex pattern suggests active metabolic compensation mechanisms that may preserve cellular membrane integrity and energy homeostasis. Progressive IPF/ILD exhibited a more constrained lipid signature with limited TG diversity and predominantly decreased abundance across multiple species. This pattern indicates compromised lipid synthetic capacity and suggests a shift toward metabolic pathways that may be insufficient to maintain cellular homeostasis during disease progression

Further, progressive disease was characterized by a relatively high abundance of amino acid metabolites, including taurine and aspartic acid, alongside increased purine metabolism indicators and hypoxanthine. Stable IPF/ILD individuals also exhibited evidence of active metabolic processes with increased betaine and acylcarnitine, suggesting ongoing methylation reactions and fatty acid oxidation, though it also shows decreased lactic acid and some acylcarnitine species, indicating complex alterations in energy metabolism.

The observed dysregulation of lipid and amino acid metabolism aligns with previous reports in IPF/ILD. These metabolic alterations reflect disrupted cellular energetics, as amino acids serve as carbon donors for tricarboxylic acid (TCA) cycle intermediates essential for fatty acid, glucose, and ATP generation^12^. For instance, the conversion of alpha-ketoglutarate to glutamate results in the synthesis of multiple amino acids, including alanine, aspartate, and arginine. Furthermore, branched-chain amino acids (BCAAs; valine, leucine, and isoleucine) can contribute to TCA cycle anaplerosis by generating acetyl-CoA^13^. Our findings are concordant with prior studies that have identified dysregulation of amino acids among other metabolites in serum and lung tissue specimens from fibrotic and non-fibrotic IPF/ILD individuals^14,15^. Specifically, a relatively high abundance of serum aspartate in progressors reinforces the potential role of amino acid metabolism in progressive disease. Our metabolomic profiling highlights taurine’s significant metabolic perturbations in the context of progressive fibrotic lung disease. Its observed high abundance is particularly compelling given its established lung-protective mechanisms, including antioxidant and anti-inflammatory actions previously characterized in bleomycin-induced fibrosis models.^16,17^ Notably, the positive correlation between purine metabolism, hypoxanthine, fatty acids and HRCT scores suggest that cellular energy metabolism dysfunction parallels the extent of radiographic abnormalities.

Purine metabolites play a crucial role in various cellular processes, including energy metabolism, signaling, and nucleotide synthesis.^18^ Moreover, the activation of the mechanistic target of rapamycin complex 1 (mTORC1) signaling pathway, which is sensitive to changes in cellular purine levels^19^, has been implicated in the pathogenesis of lung fibrosis.^20,21^ Though our study revealed low abundance of lactic acid in stable IPF/ILD individuals, there is scientific literature consistent with glycolytic shifts and increased lactate production in fibrotic lung pathologies^14,22,23^.

Sphingolipid metabolism showed dysregulation, with elevated sphingomyelins and ceramides in progressive disease. From a radiologic perspective, these findings are significant because sphingolipids serve as critical structural membrane components and precursors for pro-apoptotic and inflammatory signaling molecules implicated in fibrotic progression. Further, sphingomyelins, key structural components of cell membranes and precursors for ceramides implicated in apoptosis and pro-inflammatory signaling, were markedly elevated in both stable and progressive cohorts^24^. The observed disturbance of sphingolipids aligns with prior reports that highlight the role of ceramide and sphingolipid dysregulation in promoting fibrotic and inflammatory processes in pulmonary fibrosis.^25–27^

Lastly, predominant lipid-centric dysregulation associated with radiographic patterns of fibrosis in our study emerged as a critical hallmark of fibrotic lung disease, characterized by profound alterations in multiple lipid classes, including TGs, CEs, PI, PC, PG, LPCs, ceramides, and SMs. As we mentioned, progressors in our cohort demonstrated more strained lipid dysregulation than the stable disease cohort. There is substantial evidence of aberrant lipid metabolism in fibrotic lung disease, particularly alterations in alveolar epithelial cells (AECs) and the subsequent effects on fibroblast activation and extracellular matrix deposition.^28–31^ All in all, the observed metabolic heterogeneity between stable and progressive IPF/ILD provides critical insights into the molecular underpinnings of disease progression.

The clinical implications of our study are profound. This study is the first to correlate HRCT visual scoring with metabolomic profiling, revealing distinct metabolic patterns linked to radiological features, including ground-glass opacities, reticulation, traction bronchiectasis, and honeycombing. By systematically quantifying four cardinal radiographic features across six lung zones, the approach enables detailed, reproducible assessment of disease extent, providing a more precise phenotypic characterization of IPF/ILD. These metabolomic signatures require validation in external cohorts before their potential as predictive or diagnostic tools can be established. Future longitudinal studies will be necessary to determine whether metabolic changes precede radiographic progression. Additionally, we aim to perform targeted lipidomic, sphingomyelin, and amino acid analysis for our future research study to validate our current findings. The development of targeted therapies aimed at modulating lipids, amino acids, and purine metabolism may hold therapeutic potential for IPF/ILD patients.

Several limitations warrant consideration. The cross-sectional study design with retrospective disease progression assessment limits our ability to establish causal relationships between metabolic alterations and disease progression. Our comparison between stable and progressive groups may reflect differences in disease severity at the time of sampling rather than true progression mechanisms. Future studies should incorporate longitudinal sampling, tissue-specific metabolomic analyses, and larger cohorts to validate these findings and assess their diagnostic utility.

The metabolic differences observed between stable and progressive IPF/ILD suggest that progressive disease states are associated with heightened amino acid dysregulation (taurine), sphingomyelin, and certain triacylglycerol species. These findings signal toward more active cellular remodeling, inflammation, and energy metabolism shifts in progressive IPF/ILD compared to stable IPF/ILD.

---

**Supplementary Figure 1.**
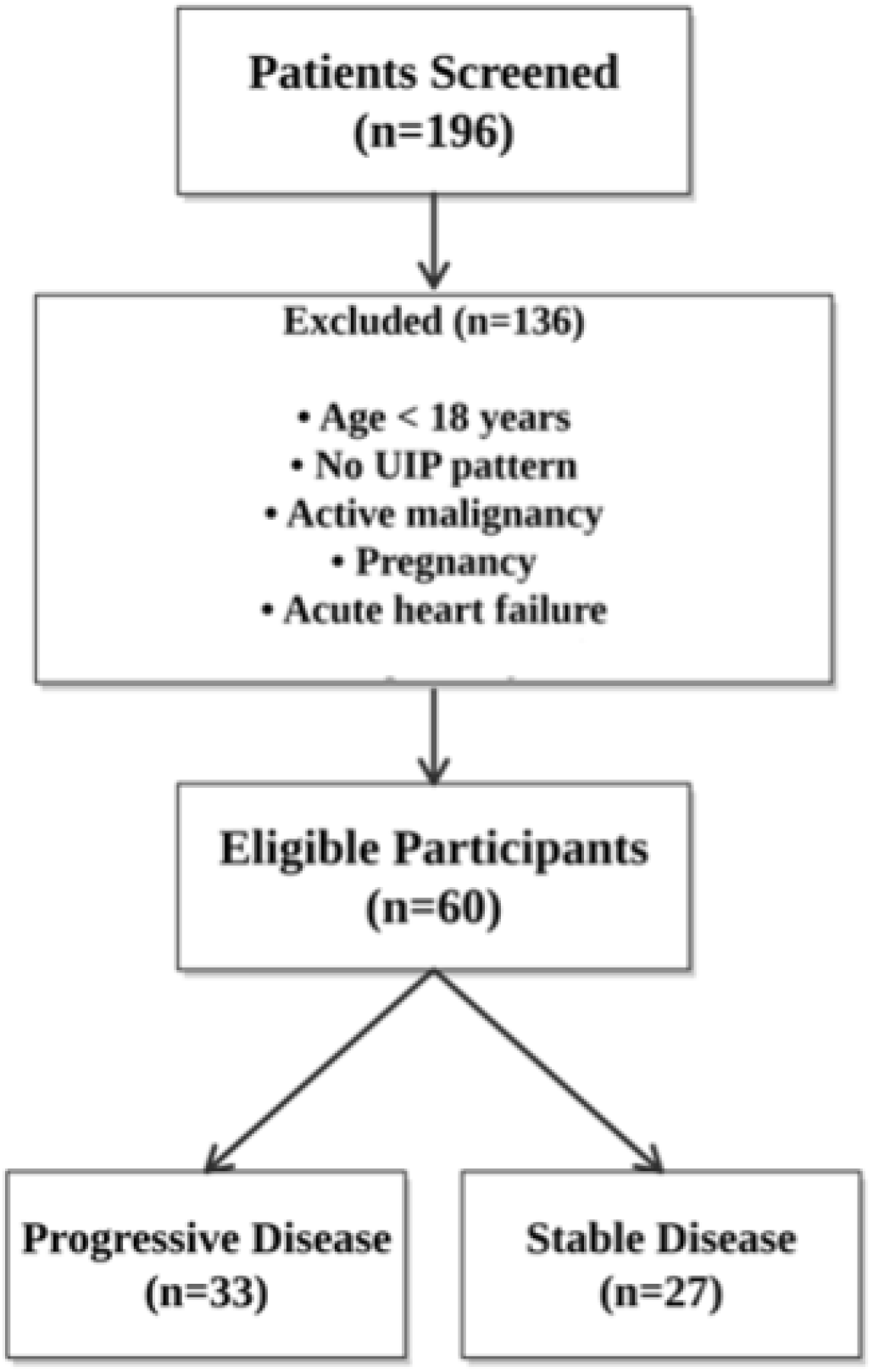
This flow diagram illustrates the patient selection process for the study. From an initial screening of 196 patients, 136 were excluded based on predefined criteria. Of the 60 eligible participants, 33 were classified as having Progressive Disease and 27 as having Stable Disease based on 2018 IPF guidelines.

## References

1. Wuyts W, Papiris S, Manali ED, et al. The Burden of Progressive Fibrosing Interstitial Lung Disease: A DELPHI Approach. Adv Ther 2020;37(7):3246–3264.

2. Raghu G, Remy-Jardin M, Richeldi L, et al. Idiopathic pulmonary fibrosis (an update) and progressive pulmonary fibrosis in adults: an official ATS/ERS/JRS/ALAT clinical practice guideline. Am J Respir Crit Care Med 2022;205(9):e18–e47.

3. Salisbury ML, Lynch DA, Beek EJR Van, et al. Idiopathic pulmonary fibrosis: the association between the adaptive multiple features method and fibrosis outcomes. Am J Respir Crit Care Med 2017;195(7):921–929.

4. Ohkubo H, Taniguchi H, Kondoh Y, et al. A volumetric computed tomography analysis of the normal lung in idiopathic pulmonary fibrosis: the relationship with the survival. Internal Medicine 2018;57(7):929–937.

5. Humphries SM, Swigris JJ, Brown KK, et al. Quantitative high-resolution computed tomography fibrosis score: performance characteristics in idiopathic pulmonary fibrosis. European Respiratory Journal 2018;52(3).

6. Taha N, D’Amato D, Hosein K, et al. Longitudinal functional changes with clinically significant radiographic progression in idiopathic pulmonary fibrosis: are we following the right parameters? Respir Res 2020;21:1–8.

7. Kazerooni EA, Martinez FJ, Flint A, et al. Thin-section CT obtained at 10-mm increments versus limited three-level thin-section CT for idiopathic pulmonary fibrosis: correlation with pathologic scoring. AJR Am J Roentgenol 1997;169(4):977–983.

8. Maldonado F, Moua T, Rajagopalan S, et al. Automated Quantification of Radiological Patterns Predicts Survival in Idiopathic Pulmonary Fibrosis. European Respiratory Journal 2013;43(1):204–212.

9. Oda K, Ishimoto H, Yatera K, et al. High-resolution CT scoring system-based grading scale predicts the clinical outcomes in patients with idiopathic pulmonary fibrosis. Respir Res 2014;15:1–9.

10. Raghu G, Remy-Jardin M, Myers JL, et al. Diagnosis of idiopathic pulmonary fibrosis. An official ATS/ERS/JRS/ALAT clinical practice guideline. Am J Respir Crit Care Med 2018;198(5):e44–e68.

11. Ravanbakhsh S, Liu P, Bjordahl TC, et al. Accurate, fully-automated NMR spectral profiling for metabolomics. PLoS One 2015;10(5):e0124219.

12. Chandel NS. Amino acid metabolism. Cold Spring Harb Perspect Biol 2021;13(4):a040584.

13. Neinast M, Murashige D, Arany Z. Branched chain amino acids. Annu Rev Physiol 2019;81(1):139–164.

14. Zhao YD, Yin L, Archer S, et al. Metabolic heterogeneity of idiopathic pulmonary fibrosis: a metabolomic study. BMJ Open Respir Res 2017;4(1):e000183.

15. Zhan J, Jarrell ZR, Hu X, et al. A pilot metabolomics study across the continuum of interstitial lung disease fibrosis severity. Physiol Rep 2024;12(20):e70093.

16. Gurujeyalakshmi G, Wang Y, Giri SN. Suppression of bleomycin-induced nitric oxide production in mice by taurine and niacin. Nitric Oxide 2000;4(4):399–411.

17. Giri SN. The combined treatment with taurine and niacin blocks the bleomycin-induced activation of nuclear factor-kB and lung fibrosis. In: Taurine 5: Beginning the 21st Century. Springer; 2003. p. 381–394.

18. Huang Z, Xie N, Illes P, et al. From purines to purinergic signalling: molecular functions and human diseases. Signal Transduct Target Ther 2021;6(1):162.

19. Hoxhaj G, Hughes-Hallett J, Timson RC, et al. The mTORC1 signaling network senses changes in cellular purine nucleotide levels. Cell Rep 2017;21(5):1331–1346.

20. Selvarajah B, Azuelos I, Platé M, et al. mTORC1 amplifies the ATF4-dependent de novo serine-glycine pathway to supply glycine during TGF-β1–induced collagen biosynthesis. Sci Signal 2019;12(582):eaav3048.

21. Woodcock H V, Eley JD, Guillotin D, et al. The mTORC1/4E-BP1 axis represents a critical signaling node during fibrogenesis. Nat Commun 2019;10(1):6.

22. Kang YP, Lee SB, Lee J, et al. Metabolic profiling regarding pathogenesis of idiopathic pulmonary fibrosis. J Proteome Res 2016;15(5):1717–1724.

23. Roque W, Romero F. Cellular metabolomics of pulmonary fibrosis, from amino acids to lipids. American Journal of Physiology-Cell Physiology 2021;320(5):C689–C695.

24. Harvald EB, Olsen ASB, Færgeman NJ. Autophagy in the light of sphingolipid metabolism. Apoptosis 2015;20:658–670.

25. Nambiar S, Clynick B, How BS, et al. There is detectable variation in the lipidomic profile between stable and progressive patients with idiopathic pulmonary fibrosis (IPF). Respir Res 2021;22:1–8.

26. Kulkarni YM, Dutta S, Iyer AK V, et al. A lipidomics approach to identifying key lipid species involved in VEGF-inhibitor mediated attenuation of bleomycin-induced pulmonary fibrosis. PROTEOMICS–Clinical Applications 2018;12(3):1700086.

27. Jayant G, Kuperberg S, Somnay K, Wadgaonkar R. The role of sphingolipids in regulating vascular permeability in idiopathic pulmonary fibrosis. Biomedicines 2023;11(6):1728.

28. Yang J, Pan X, Xu M, et al. Downregulation of HMGCS2 mediated AECIIs lipid metabolic alteration promotes pulmonary fibrosis by activating fibroblasts. Respir Res 2024;25(1):176.

29. Shi X, Chen Y, Shi M, et al. The novel molecular mechanism of pulmonary fibrosis: insight into lipid metabolism from reanalysis of single-cell RNA-seq databases. Lipids Health Dis 2024;23(1):98.

30. Agudelo CW, Samaha G, Garcia-Arcos I. Alveolar lipids in pulmonary disease. A review. Lipids Health Dis 2020;19(1):122.

31. O’Callaghan M, Tarling EJ, Bridges JP, et al. Reexamining the Role of Pulmonary Lipids in the Pathogenesis of Pulmonary Fibrosis. Am J Respir Cell Mol Biol 2024;71(4):407–419.

